# Production of nonnatural straight-chain amino acid 6-aminocaproate via an artificial iterative carbon-chain-extension cycle

**DOI:** 10.1101/568121

**Authors:** Jie Cheng, Tingting Song, Huayu Wang, Xiaohua Zhou, Michael P. Torrens-Spence, Dan Wang, Jing-Ke Weng, Qinhong Wang

**Affiliations:** Department of Chemical Engineering, School of Chemistry and Chemical Engineering, Chongqing University, Chongqing 401331, P. R. China; Key Laboratory of Systems Microbial Biotechnology, Tianjin Institute of Industrial Biotechnology, Chinese Academy of Sciences, Tianjin 300308, P. R. China; Whitehead Institute for Biomedical Research, 455 Main Street, Cambridge, MA, 02142, United States; Department of Biology, Massachusetts Institute of Technology, Cambridge, MA, United States

**Keywords:** Nonnatural straight chain amino acid, 6-Aminocaproate, Carbon chain elongation, Synthetic biology, Iterative cycle

## Abstract

Bioplastics produced from microbial source are promising green alternatives to traditional petrochemical-derived plastics. Nonnatural straight-chain amino acids, especially 5-aminovalerate, 6-aminocaproate and 7-aminoheptanoate are potential monomers for the synthesis of polymeric bioplastics as their primary amine and carboxylic acid are ideal functional groups for polymerization. Previous pathways for 5-aminovalerate and 6-aminocaproate biosynthesis in microorganisms are derived from L-lysine catabolism and citric acid cycle, respectively. Here, we show the construction of an artificial iterative carbon-chain-extension cycle in *Escherichia coli* for simultaneous production of a series of nonnatural amino acids with varying chain length. Overexpression of L-lysine α-oxidase in *E. coli* yields 2-keto-6-aminocaproate as a non-native substrate for the artificial iterative carbon-chain-extension cycle. The chain-extended α-ketoacid is subsequently decarboxylated and oxidized by an α-ketoacid decarboxylase and an aldehyde dehydrogenase, respectively, to yield the nonnatural straight-chain amino acid products. The engineered system demonstrated simultaneous *in vitro* production of 99.16 mg/L of 5-aminovalerate, 46.96 mg/L of 6-aminocaproate and 4.78 mg/L of 7-aminoheptanoate after 8 hours of enzyme catalysis starting from 2-keto-6-aminocaproate as the substrate. Furthermore, simultaneous production of 2.15 g/L of 5-aminovalerate, 24.12 mg/L of 6-aminocaproate and 4.74 mg/L of 7-aminoheptanoate was achieved in engineered *E. coli*. This work illustrates a promising metabolic-engineering strategy to access other medium-chain organic acids with -NH_2_,-SCH_3_, -SOCH_3_, -SH, -COOH, -COH, or -OH functional groups through carbon-chain-elongation chemistry.

## 1. Introduction

Microbial polyimide bioplastics present a class of green materials with broad applications in many downstream industries, and can potentially replace the traditional petrochemical-derived polymers. Consequently, platform chemicals containing suitable functional groups necessary for polyimide polymerization have attracted significant attention as targets for metabolic engineering. These compounds include diamines such as putrescine (Del Rio et al., 2018) and cadaverine (Kim et al., 2018), amino acids such as lysine (Borri et al., 2018) and glutamate, organic acids such as succinate (Jantama et al., 2008) and lactate (Pang et al., 2010), diols such as butanediol and hexanediol (Muller et al., 2010). Nonnatural straight-chain amino acids (NNSCAAs), especially 5-aminovalerate (5AVA) and 6-aminocaproate (6ACA) are important platform chemicals for the synthesis of polyimides, which are widely used as raw materials for automobile parts, clothes, backpacks and disposable goods such as nylon 5, nylon 6 and nylon 5,6 (Haushalter et al., 2017). In addition to its utility in bioplastics, 6ACA was also implicated to promote blood clotting, suggesting potential applications as an antifibrinolytic agent (Lu et al., 2015; Schou-Pedersen et al., 2015). Whereas 5AVA biosynthesis is a viable approach for industrial production, effective methods to biosynthesize other NNSCAAs at scale has yet to be established (Jorge et al., 2017; Turk et al., 2016). Biosynthesis of 6ACA was first demonstrated to occur through the condensation of acetyl-CoA and succinyl-CoA (Turk et al., 2016). The second biosynthetic route utilizes α-ketoadipate as the starter molecule, which is chain-extended by (homo)_1 →3_aconitate synthase (AksA), (homo)_1 →3_aconitate isomerase complex (AksD, AksE), iso(homo)_1 →3_citrate dehydrogenase (AksF) to give the intermediate α-ketopimelate (AKP). AKP is decarboxylated and transaminased to yield 6ACA (Chae et al., 2017). The precursors of the two pathways are all derived from the tricarboxylic acid (TCA) cycle which are scarce in cells. With inadequate transamination efficiency previously recognized (Zhang et al., 2010), the final titer achieved by Turk *et al.* was only 160 mg/L (Jorge et al., 2017; Turk et al., 2016).

L-lysine is the second most-produced amino acid worldwide after glutamate. Currently, L-lysine is mainly produced through microbial fermentation, and is commonly used as an additive to poultry and swine feed (Wang et al., 2016). Annual world L-lysine production is estimated to exceed 2.5 million tons by 2020 (Vassilev et al., 2018). Due to the market competition in industrial capacity and demand, the price of L-lysine as a commodity chemical has dropped significantly in recent years (Xu et al., 2018). As a result, developing high-value chemicals derived from L-lysine presents an emerging opportunity in the field of metabolic engineering (Cheng et al., 2018a). Novel L-lysine-derived products may contribute to an environmentally friendly chemical industry (Hoffmann et al., 2018; Sgobba et al., 2018).

Nonpolymeric carbon-chain-extension pathways occur broadly in many primary metabolism pathways for the synthesis of rare sugars (Yang et al., 2017), α-ketoacids (Sonderby et al., 2010; Wen et al., 2013), fatty acids (Wu et al., 2014), and as well as several specialized metabolic pathways for the synthesis of polyketides (Gokhale et al., 2007; Miyanaga, 2017) and terpenoids (Gronenberg et al., 2013; Yu et al., 2012). The chain extension of the aforementioned metabolic systems generally consists of a series of condensation, dehydration and reduction reactions (Chandran et al., 2011; Katz and Donadio, 1993; Textor et al., 2007). The first carbon-chain-extension step of the pathway is catalyzed by a C-acetyltransferase, such as the citramalate synthase in the citramalate pathway (Drevland et al., 2007), the citrate synthase in the TCA (Harder et al., 2018), the α-isopropylmalate synthase (LeuA) in the leucine pathway (Hunter and Parker, 2014), the homocitrate synthase in the L-lysine α-aminoadipate pathway (Zabriskie and Jackson, 2000), the methylthioalkylmalate synthase (MAM) in the glucosinolate pathway (Mirza et al., 2016) and the (Homo)_1→3_aconitate synthase (AksA) in the biosynthetic pathway of coenzyme B (Howell et al., 1998). Amongst various C-acetyltransferases, LeuA is paid much more attention. In its native pathway, LeuA catalyzes the condensation of acetyl-CoA and α-ketoisovalerate in the first step of leucine biosynthesis (Shen and Liao, 2011). LeuA could also catalyze the condensation of acety-CoA and α-ketobutyrate to produce α-ketovalerate as a precursor of the non-natural amino acid (Shen and Liao, 2008). Overexpression of LeuABCD, α-ketoacid decarboxylase (KivD) and alcohol dehydrogenase (ADH6) in a modified threonine-overproduction strain of *Escherichia coli* (*E. coli*) ATCC98082/ThrABC-TdcB-IlvGMCD led to the production of (*S*)-3-methyl-1-pentanol with a final titer of 6.5 mg/L (Zhang et al., 2008). However, several mutants derived from LeuA exhibit interesting substrate promiscuity and catalyzes flexibility, which we use LeuA* to refer to (Shen and Liao, 2008; Umbarger, 1978). LeuA* displays an enhanced degree of substrate promiscuity and is capable of catalyzing the condensation reaction on multiple α-ketoacid substrates (Shen and Liao, 2008). A mutant with LeuA* G462D to replace the LeuA in engineered strain ATCC98082/ThrABC-TdcB-IlvGMCD-LeuA*BCD-KIVD-ADH6 led to an improved titer of (*S*)-3-methyl-1-pentanol at 40.8 mg/L. Another LeuA* G462D/S139G mutant could improve the titer of 1-pentanol to 204.7 mg/L (Zhang et al., 2008). Nevertheless, only limited functional groups have been successfully introduced into ketoacid elongation pathway.

In this study, we explore the use of an L-lysine-derived α-ketoacid with -NH_2_functional group as a substrate for LeuABCD-catalyzed carbon-chain-extensions. We build an artificial iterative carbon-chain-extension cycle for NNSCAA biosynthesis from L-lysine as seen in Fig. 1. We demonstrate that NNSCAAs of C5, C6 and C7 could be simultaneously produced in engineered *E. coli* strain.

**Fig. 1.**
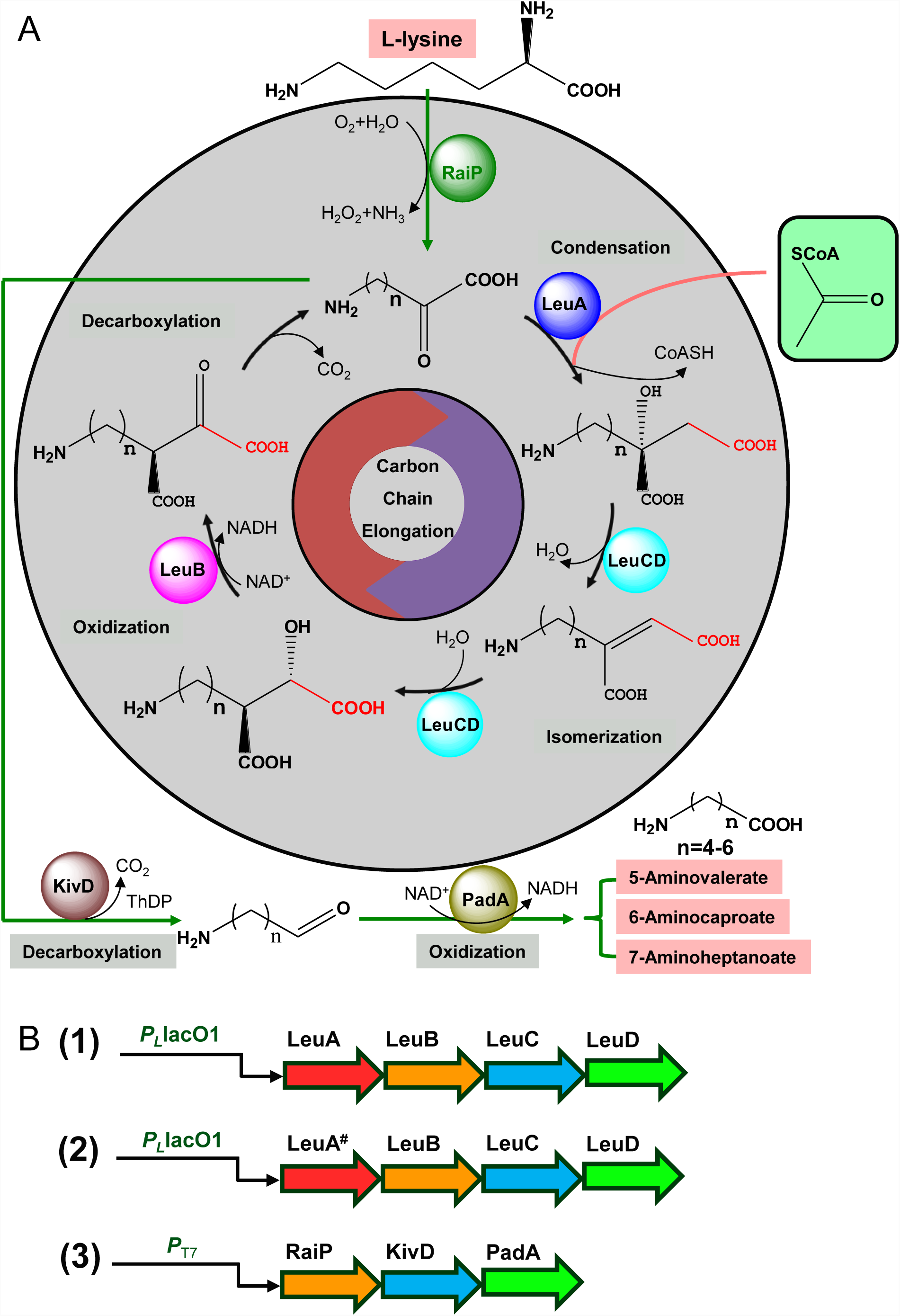
A: Engineered artificial iterative carbon-chain-extension cycle for the production of NNSCAAs. B: Synthetic operons for gene expression. (1) Synthetic operon for protein overexpression to drive the carbon flux towards 2-keto-6-aminocaproate. (2) Synthetic operon carrying mutations of LeuA (LeuA^#^) for protein overexpression to drive the carbon flux towards 2-keto-6-aminocaproate. (3) Synthetic operon for protein overexpression for deaminase, decarboxylase and dehydrogenase. NNSCAAs, Nonnatural straight chain amino acids. RaiP, L-lysine α oxidase; LeuA, α-Isopropylmalate synthase; LeuA*, LeuA mutants; LeuA^#^, LeuA with H97L/S139G/G462D mutations; LeuB, 3-Isopropylmalate dehydrogenase; LeuC, 3-Isopropylmalate dehydratase; LeuD, 3-Isopropylmalate dehydratase; KivD, α-Ketoacid decarboxylase; PadA, Phenylacetaldehyde dehydrogenase; ThDP, Thiamine diphosphate.

## 2. Material and methods

### 2.1 Strains, plasmids and primers used in this study

All the strains and plasmids involved are listed in Table S1. Primers used in this study are listed in Table S2. *E. coli* BL21(DE3) was used for the production of NNSCAAs. *E. coli* DH5α was used for plasmid amplification. The L-lysine α oxidase gene (*raiP*) from *Scomber japonicas* (GenBank Accession No. MG423617) was amplified from pCJ01 (Cheng et al., 2018b). The coding regions of *leuA*, 3-isopropylmalate dehydrogenase gene (*leuB*), 3-isopropylmalate dehydratase gene (*leuC*), 3-isopropylmalate dehydratase gene (*leuD*) and aldehyde dehydrogenase gene (*padA*) were amplified from *E. coli* MG1655 by PCR using appropriate primers (Table S2). The α-ketoacid decarboxylase gene (*kivD*) from *Lactococcus lactis* (GenBank Accession No. 51870501) was chemically and optimally synthesized by Genewiz Co. (Suzhou, China) (Li et al., 2011). In order to establish and validate the whole biosynthetic pathway, the *leuA* and *leuBCD* genes from *E. coli* MG1655 were constructed in a single operon in transcriptional order *leuA*-*leuB*-*leuC*-*leuD*, and then the engineered pZA22-*leuA*-*leuB*-*leuC*-*leuD* was produced, also named as pIVC03. LeuA was replaced by LeuA^#^(LeuA*, LeuA with H97L/S139G/G462D mutations) to form the engineered pZA22-*leuA*^#^-*leuB*-*leuC*-*leuD*, also named as pIVC04. The *raiP* from *Scomber japonicus, kivD* from *Lactococcus lactis* and *padA* from *E. coli* MG1655 were constructed in another operon in transcriptional order *raiP*-*kivD*-*padA*. The engineered pET21a-*raiP*-*kivD*-*padA* was produced, also named as pETaRKP. BL21(DE3) was transformed with the plasmid pIVC03 or pIVC04 and pETaRKP, resulting in strain CJ03 or CJ04.

### 2.2 Fermentation procedures

The fermentation media were developed for evaluating strain’s potential for NNSCAAs production. The medium was supplemented by 10 g/L NaCl, 10 g/L tryptone, 5 g/L yeast extract, 1.0 mM MgSO_4_, 0.5 mM thiamine diphosphate (ThDP) (Chen et al., 2017). For NNSCAAs production, a single colony of the desired strain was cultivated for 12 h at 37 °C and 250 RPM in 2 mL LB medium supplemented with appropriate antibiotics. This starter culture were then transferred into 40 mL of fermentation medium supplemented with appropriate antibiotics, 1.0 mM MgSO_4_and 0.5 mM ThDP at 37 °C with 2 50 RPM orbital shaking at a starting optical density at 600 nm (OD_600_) of 0.1 in a 250 mL flask. After an OD_600_of 0.6 has been reached, 0.5 mM of IPTG and 5 g/L of L-lysine were added. Flasks were then incubated at 30 °C.

### 2.3 Protein expression and purification

BL21(DE3) harboring pETaraiP, pETaleuA, pETaleuB, pETaleuC, pETaleuD, pETbkivD, pETapadA and various mutants of LeuA were screened on selective LB agar plates supplemented with 100 μg/mL ampicillin, respectively. A positive clone was inoculated in 2 mL LB at 37 °C and 250 RPM for 12 h. 2 mL of seed cultures were transferred into 200 mL LB containing 100 μg/mL ampicillin. At an OD_600_of 0.6, the cells were induced by 0.5 mM IPTG and incubated at 20 °C and 250 RPM for 16 h. The cells were collected by centrifugation at 10,000 RPM for 5 min and washed twice with potassium phosphate buffer (KPB, 50 mM, pH 8.0). The cells were resuspended and disrupted by sonication in 50 mM KPB and 2 mM tris (2-carboxyethyl) phosphine (TCEP) in an ice bath. The enzymes were purified by AKTA Purifier 10 using a Ni-NTA column (GE Healthcare, USA). The purified enzymes were desalted and exchanged into storage buffer (50 mM KPB, 1.0 mM MgSO4, 2 mM TCEP, 10% glycerol, pH 8.0) (Fan et al., 2018). The purified enzymes were stored at -80 °C. The UV absorbance at 280 nm was mensurated as the protein concentration by SpectraMax M2^e^(Molecular Devices, American) (Annamalai et al., 2011; Zhang et al., 2008). Sodium dodecyl sulfate polyacrylamide gel electrophoresis (SDS-PAGE) was used to analyze the purity of purified enzymes with a 12 % acrylamide gel (Tani et al., 2015a)2

### 2.4 Enzyme assay

RaiP activity was surveyed by determining hydrogen peroxide produced as Cheng *et al.* reported (Cheng et al., 2018b). The reaction buffer embodied 50 mM KPB (pH 8.0), 30.0 mM L-lysine, 0.5 mM 4-aminoantipyrine (4AAP), 10 units/ml catalase and 26.5 mM phenol (Muramatsu et al., 2006). The standard reaction mixture embodied 8 μg RaiP and 180 μL reaction buffer. The reaction was conducted at 30 °C and stopped by adding 10 μL of 10 M HCl. 10 μL of 10 M NaOH was added for neutralization of the reaction, and then Quinoneimine dye formed was measured at 505 nm per 5 minutes using SpectraMax M2^e^(Molecular Devices, American) (Job et al., 2002; Tani et al., 2015a; Tani et al., 2015b). One unit of enzyme activity was defined as the amount of enzyme that catalyzes 1 μmol of hydrogen peroxide produced per minute.

LeuA and LeuA* activities were assayed by measuring CoASH produced (Zhang et al., 2008). The assay mixture contained 50 mM KPB (pH 8.0), 0.03 μM acetyl-CoA, 30.0 mM L-lysine, 2.0 μM RaiP and 8 μg LeuA or LeuA* with a total volume of 205 μL. The reaction was performed at 37 °C and stopped by adding 10 μL of 10 M HCl. 10 μL of 10 M NaOH was added for neutralization of the reaction, then 50 μL of a fresh 3.0 mM solution of 5,5′-Dithio-Bis (2 Nitrobenzoic Acid) (DTNB) in 50 mM KPB was added, and the yellow color product was determined at 412 nm using SpectraMax M2^e^(Molecular Devices, American). One unit of enzyme activity was defined as the amount of enzyme that catalyzes 1.0 μmol of CoASH produced per minute. There were no other intermediates as substrates, so the enzyme activities of LeuB, LeuC, LeuD, KivD and PadA could not be detected.

### 2.5 *In vitro* 6ACA synthesis

To measure the rate of the conversion of L-lysine into 6ACA by purified enzyme, an assay mixture was established, which contained 800 μL of 50 mM KPB (pH 8.0), mM L-lysine, 2.0 mM acetyl-CoA, 0.5 mM ThDP, 1.0 mM NAD^+^, 1.0 mM TCEP. All assays were started with the addition of the different purified enzyme dosage and incubating at 37 °C. 400 μL samples were withdrawn at designated time points and inactivated at 75 °C for 10 min, and then centrifuged at 10000 RPM for 10 min to remove cells for further metabolite analysis.

### 2.6 Analytical methods

The optical density of the various *E. coli* cultures was detected using a UV/visible spectrophotometer (Ultrospec TM 2100 pro, GE Healthcare, UK). The quantification of L-lysine, 5AVA, 6ACA and 7AHA were conducted by high performance liquid chromatography (HPLC) using a 1260 system (Agilent Co., Ltd, CA, USA) with an Agilent Eclipse XDB-C18 column (4.6 mm × 150 mm × 5 μm). For detection of L-lysine, 5AVA, 6ACA and 7AHA, 360 μL of culture centrifuged was derived with phenyl isothiocyanate (PITC) (Cheng et al., 2018b; Zheng et al., 2015). The operating conditions were performed as 1.0 mL/min, column temperature 40 °C, wavelength 254 nm and analysis time 55 min. The gradient program was shown in Table S3. For liquid chromatography-mass spectrometry (LC-MS) identification of 5AVA, 6ACA and 7AHA, exact mass spectra were explored with a Bruker micrOTOF-Q II mass spectrometer using the time of flight (TOF) technique, equipped with an ESI source operating in negative mode (Burker Co., Ltd, USA). The product was verified by LC-MS (Fig. 3). The approximate retention times of 5AVA, 6ACA, 7-aminoheptanoate (7AHA) and L-lysine were 7.2 min, 9.4 min, 11.8 min and 25.6 min, respectively (Fig. 3A). LC-MS results of the 5AVA, 6ACA and 7AHA showed that the [M-H]^-^of 5AVA, 6ACA and 7AHA were 251.0801 (Fig. 3B), 265.1012 (Fig. 3C) and 279.1175 (Fig. 3D), respectively, which were as the same as that of the 5AVA, 6ACA and 7AHA standards.

## 3. Results and discussion

### 3.1 Construction of a L-lysine derived artificial iterative carbon-chain-extension cycle *in vitro*

To explore the feasibility of a RaiP-LeuABCD-KivD-PadA pathway, the necessary enzymes were expressed, purified and assayed against L-lysine for NNSCAAS synthesis. The purity of recombinant RaiP, LeuA, LeuA*, LeuB, LeuC, LeuD, KivD and PadA carrying a C-terminal His-tag in *E. coli* BL21(DE3) was assessed by SDS– PAGE shown in Fig. S1. The sizes of the recombinant proteins were 55, 52, 38, 50, 21, 57 and 52 kDa respectively, which were consistent with the predicted size of RaiP, LeuA, LeuB, LeuC, LeuD, KivD and PadA proteins. The SDS-PAGE analysis of various LeuA mutants were shown in Fig. S3.

The activities of various LeuA mutants (LeuA*) were shown in Fig. 2. LeuA exhibited low activity toward 2-keto-6-aminocaproate, whereas LeuA mutations (H97A/G462D, H97G/G462D, H97L/G462D, S139G/G462D and S139I/G462D) displayed higher activities. The LeuA H97L/S139G/G462D (LeuA^#^) showed highest activity shown in Table 1. The activities of RaiP toward L-lysine and LeuA^#^toward 2-keto-6-aminocaproate in the crude extracts were 5.14 and 0.0012 units/mg, respectively. 141-fold purification factor and 58.62% yields of RaiP were obtained, with specific activity of 724.88 units/mg shown in Table 1. 25-fold purification factor and 13.33% yields of LeuA^#^were achieved, with specific activity of 0.03 units/mg. RaiP showed a K_m_value of 2.945 mM, a K_cat_value of 710.587 s^-1^ and a K_cat_/K_m_value of 241292 M^-1^s^-1^when L-lysine is used as the substrate. LeuA^#^ displays a K_m_value of 23.450 mM, a K_cat_value of 1.321 s^-1^ and a K_cat_/K_m_value of 26.344 M^-1^s^-1^ with 2-keto-6-aminocaproate used as the substrate shown in Table 2.

**Table 1.** Purification of L-lysine α oxidase (RaiP) from *Scomber japonicas* and α-isopropylmalate synthase mutant LeuA# (LeuA with H97L/S139G/G462D mutations) expressed in *E. coli*.

| Enzyme | Step | Total protein (mg) | Total activity (units) | Specific activity (units/mg) | Purification (fold) | Yield (%) |
| --- | --- | --- | --- | --- | --- | --- |
| RaiP | Cell extract | 310.23±21.84 | 1595.21±78.52 | 5.14±0.26 | 1 | 100 |
| | Ni-NTA | 1.29±0.09 | 935.09±65.48 | 724.88±46.24 | 141 | 58.62 |
| LeuA<sup>#</sup> | Cell extract | 521.51±32.16 | 0.60±0.01 | 0.0012±0.0001 | 1 | 100 |
| | Ni-NTA | 2.97±0.13 | 0.08±0.01 | 0.03±0.002 | 25 | 13.33 |

Data are presented as means ± STDV calculated from at least three replicates. The RaiP activity was conducted on 50 mM KPB (pH 8.0), 30.0 mM L-lysine, 0.5 mM 4-aminoantipyrine, 10 units/ml catalase and 26.5 mM phenol, 8 μg RaiP with a total volume of 205 μL. The reaction was incubated at 30 °C and stopped by adding 10 μL of 10 M HCl. The quinoneimine dye formed was measured at 505 nm per 5 minutes. The LeuA<sup>#</sup> activity was performed on 50 mM KPB (pH 8.0), 0.03 μM acetyl-CoA, 30.0 mM L-lysine, 2.0 μM RaiP and 8 μg LeuA<sup>#</sup> with a total volume of 205 μL. The reaction was performed at 37 °C and stopped by adding 10 μL of 10 M HCl. 50 μL of a fresh 3.0 mM solution of DTNB in 50 mM KPB was added, and the yellow color product was determined at 412 nm.

**Table 2.**
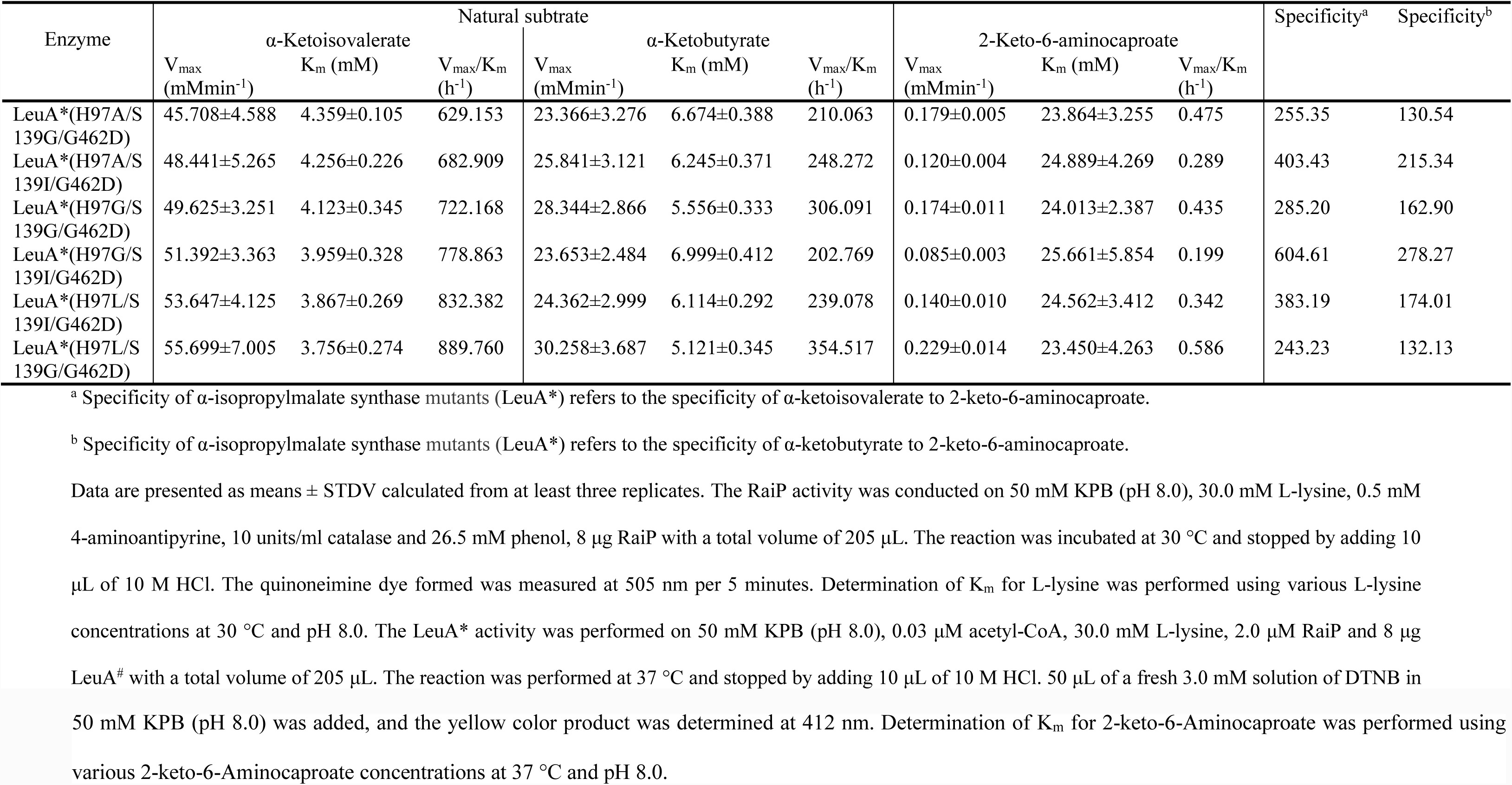
Kinetic parameters of α-isopropylmalate synthase mutants (LeuA*) on 2-keto-6-aminocaproate.

**Fig. 2.**
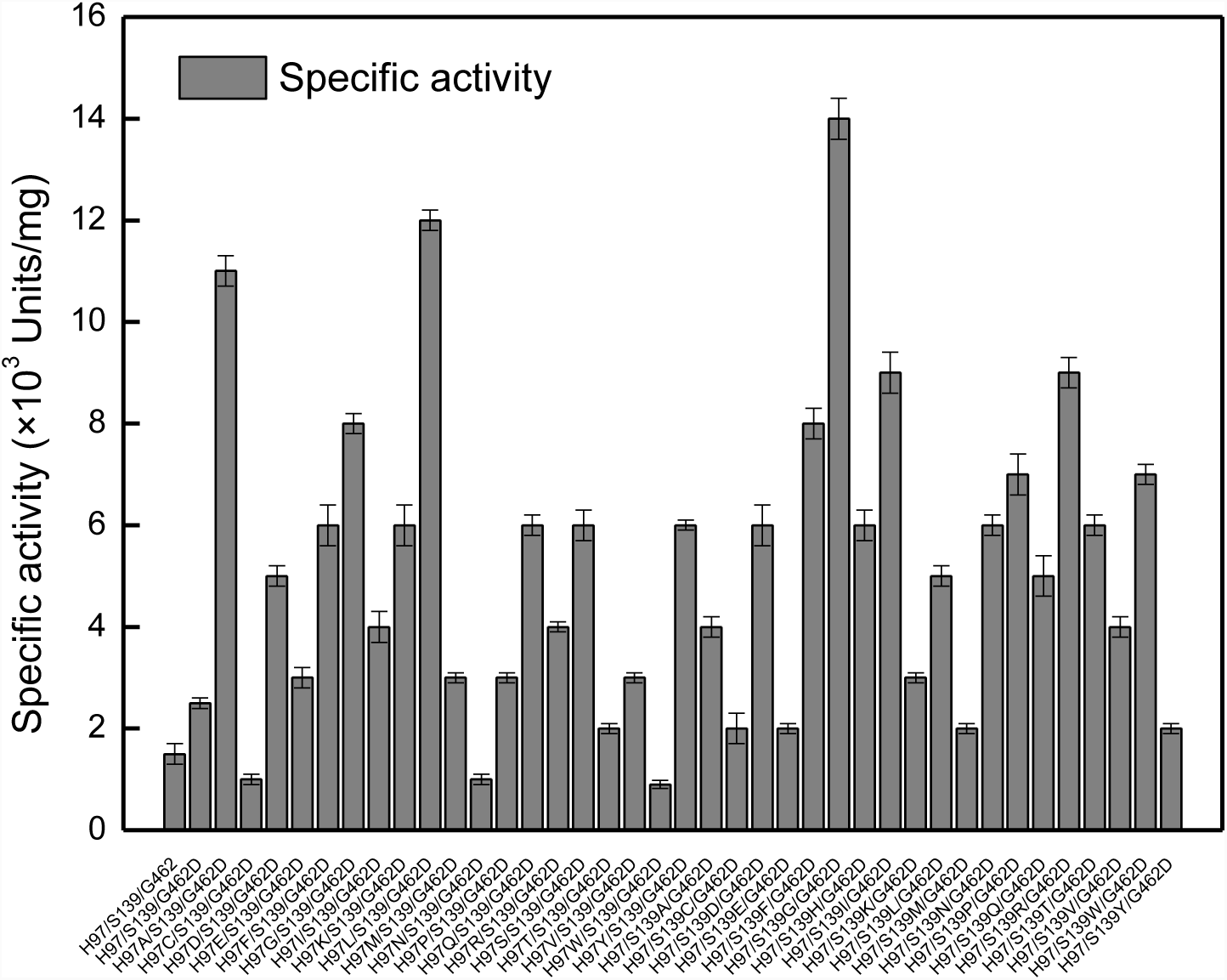
Specific activities of various LeuA mutants. The assay mixture contained 50 mM KPB, 0.03 μM acetyl-CoA and 30.0 mM L-lysine, 2.0 μM RaiP and 8 μg LeuA* with a total volume of 205 μL. All experiments were performed a minimum of three independent sets. All error bars represent standard deviations with n ≥ 3 independent reactions.

**Fig. 3.**
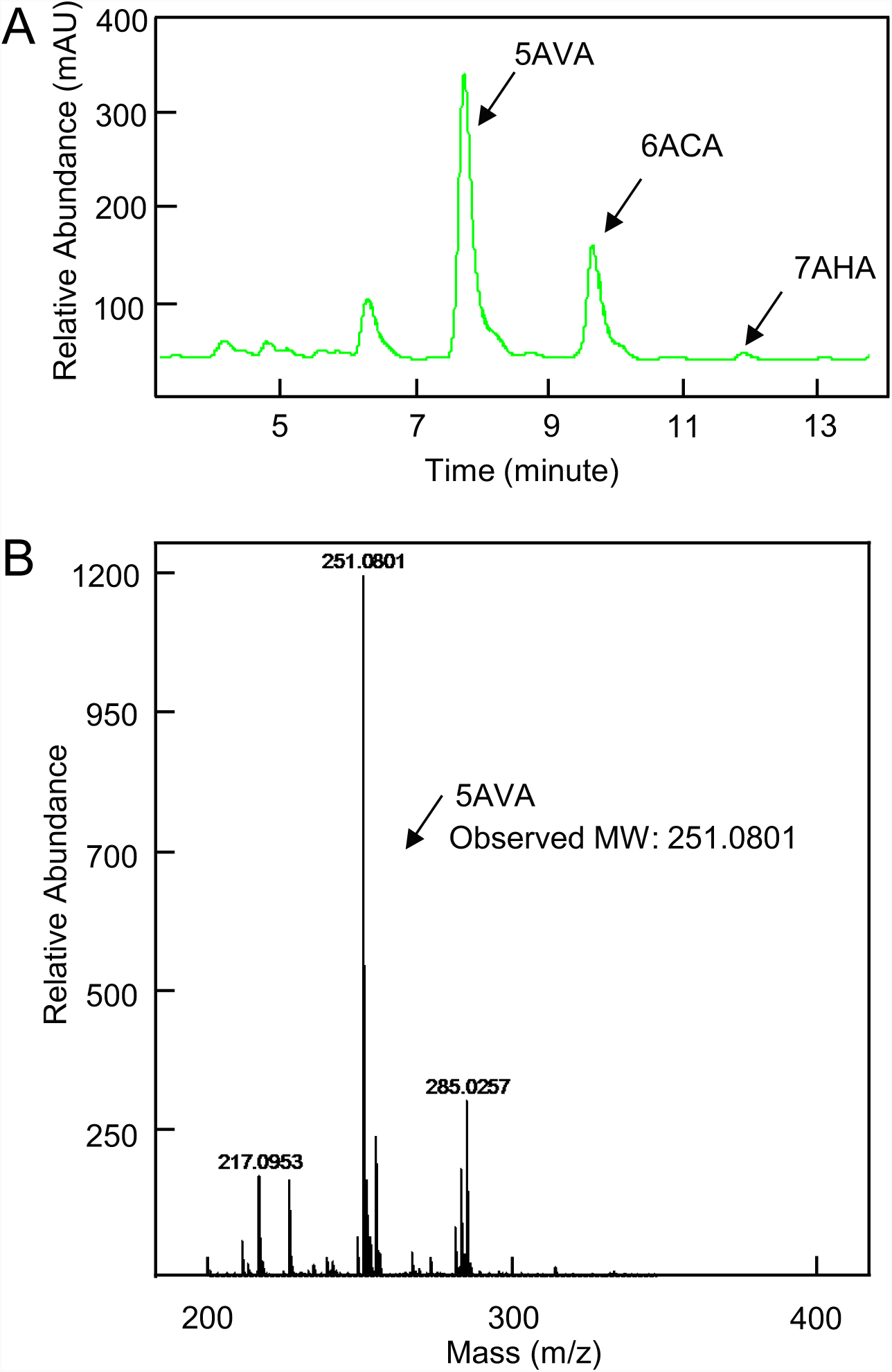
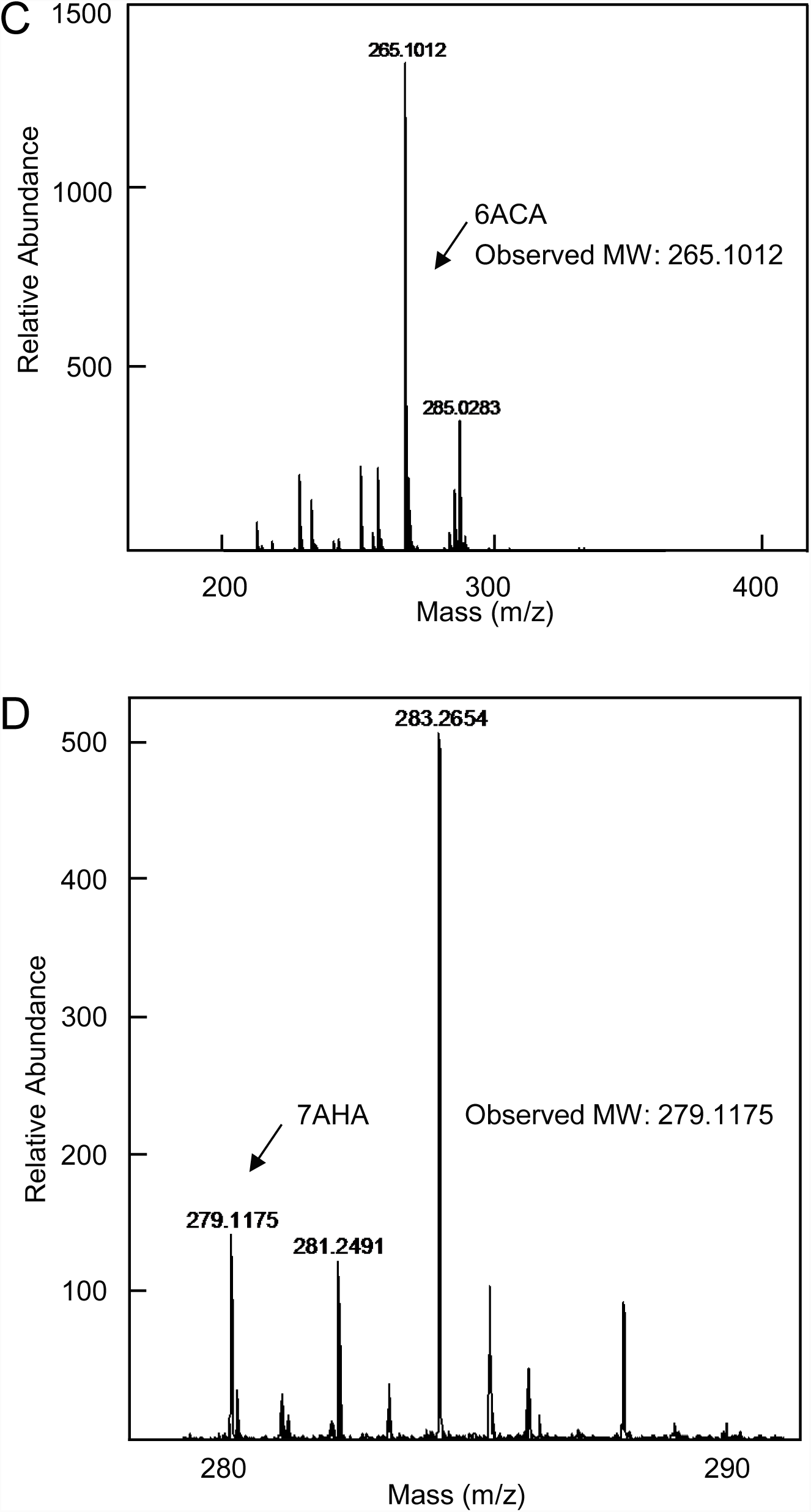
LC-MS confirmation of NNSCAAs (5AVA, 6ACA, 7AHA) biosynthesis by strain CJ04. A: HPLC results of 5AVA, 6ACA and 7AHA from fermentation broth. B: Mass spectrum of 5AVA from fermentation broth. C: Mass spectrum of 6ACA from fermentation broth. D: Mass spectrum of 7AHA from fermentation broth. Samples were derived with phenyl isothiocyanate (PITC) for LC-MS analysis. Strain CJ04 is the strain BL21(DE3) harboring plasmids pIVC4 and pET21aRKP. 5AVA, 5-Aminovalerate. 6ACA, 6-Aminocaproate. 7AHA, 7-Aminoheptanoate.

Amano *et al.* and Arinbasarova *et al.* have previously characterized the recombinant enzyme of RaiP from *Trichoderma viride* (Amano et al., 2015; Arinbasarova et al., 2012). However, the reported specific activities of the purified enzyme was just 80 or 90 U/mg in the previous studies, which are about 11% of the specific activity we measured in this study. It is likely that a fraction of the enzyme might have lost its activity in the process of ammonium sulfate precipitation and two-step column purification in the previous studies (Tani et al., 2015b). Zhang *et al.* reported that the LeuA G462D mutant displays a low K_cat_of 0.018 s^-1^ for (*S*)-2-keto-3-methylvalerate (Zhang et al., 2008). However, the LeuA S139G/G462D mutant exhibits a much higher K_cat_of 0.12 s^-1^ for (*S*)-2-keto-3-methylvalerate. We therefore introduced an additional mutations at His97 in LeuA^#^. Interestingly, LeuA^#^shows significantly improved K_cat_of 1.32 s^-1^ for 2-keto-6-aminocaproate shown in Table 2.

### 3.2 Building a nonnatural iterative cycle for NNSCAA biosynthesis *in vitro*

Upon the successful engineering of LeuA, we designed a novel NNSCAA biosynthetic pathway by combining a “+1” carbon-chain-extension pathway with subsequent α-ketoacid decarboxylation and oxidation. The promiscuity of various LeuA mutants was explored, which resulted in an iterative cycle for the production of a series of L-lysine-derived products with different chain lengths (Fig. 6A and Fig. 6B). The titer of total NNSCAAs was 65.92 mg/L after a reaction time of 4 h with enzyme concentration of RaiP, LeuA^#^, LeuB, LeuC, LeuD, KivD and PadA setting at 1.0, 20.0, 4.0, 2.0, 2.0, 5.0 and 2.0 μM, respectively. The distribution of 5AVA, 6ACA and 7AHA was 24.0:5.2:1. As could be seen in Fig. 6, with a prolonged reaction time of 8 h, the titer of total NNSCAAs reached 151.28 mg/L, and the distribution of 5AVA, 6ACA and 7AHA was 19.2:9.1:1. The identity of the products were verified by LC-MS as shown in Fig. 3.

NNSCAAs, especially 5AVA, 6ACA and 7AHA, are important platform chemicals for polyamides synthesis. To our knowledge, simultaneous production of NNSCAAs of various chain length has not been demonstrated in any microbial host. Specific pathways for the production of 5AVA or 6ACA have been developed recently (Cheng et al., 2018b; Turk et al., 2016). The precursors are derived from L-lysine catabolism and TCA cycle, respectively. The engineered *Corynebacterium glutamicum* strain expressing *deta*-aminovaleramidase gene as an operon under a synthetic H_36_promoter reportedly produces 5AVA at 33.1 g/L (Shin et al., 2016). 5AVA could be obtained at a higher titer of 63.2 g/L by overexpressing an L-lysine-specific permease (Li et al., 2016). As the transamination activity is limiting (Zhang et al., 2010), the final titer of 6ACA achieved in Turk *et al.*’s study was only 160 mg/L (Jorge et al., 2017; Turk et al., 2016). In our work, we aimed to develop a strategy to simultaneously produce C5, C6 and C7 NNSCAAs from L-lysine. To do this, we explored the promiscuity of LeuA mutants towards L-lysine-derived α-ketoacids with amino functional group, which is exemplified by LeuA^#^that can utilize primary amines such as 2-keto-6-aminocaproate and 2-keto-7-aminoheptanoate as substrate. The malleability of the LeuABCD pathway remains to be further explored, as untargeted metabolomics of LeuA* expression *in vivo* may identify additional substrates. Furthermore, directed evolution of LeuA or LeuA* may further broaden substrate profile. In *Brassicaceae* plants, Methylthioalkylmalate synthases are evolutionary derived from an ancestral LeuA and catalyze carbon-chain-extension pathway in the biosynthesis of glucosinolates (de Kraker and Gershenzon, 2011; Mirza et al., 2016). The recruitment of LeuA for plant specialized metabolism suggests that the C-acetyltransferase family proteins can be further evolved to arrive at desirable activities starting from ancestral promiscuous activities (Weng and Noel, 2012).

### 3.2 Dependence of 6ACA productivity on the supply of coenzyme

In this artificial iterative cycle pathway, NAD^+^is a key coenzyme for LeuB. Moreover, KivD and PadA catalyze the conversion of 2-keto-7-aminoheptanoate to 6ACA, which requires coenzymes ThDP and NAD^+^, respectively (Fig. 1). To improve the production of 6ACA, the concentration of NAD^+^was optimized. Fig. 5A shows the production of 6ACA with varying concentrations of NAD^+^at pH 8.0 for 8 h. NAD^+^concentration affects the production of 6ACA shown in Fig. 5A. This multi-enzyme cascade system requires NAD^+^addition to produce 6ACA. When 0.2 mM NAD^+^was added, the titer of 6ACA was 21.24 mg/L. Increasing concentration of NAD^+^led to increase titer of 6ACA production. Notably, the concentration of 6ACA increased to 46.96 mg/L with 1 mM NAD^+^. Hence, 1 mM of NAD^+^was set as the optimal dosage. The effect of ThDP, the coenzyme of KivD, was also investigated in this work. The results were shown in Fig. 5B. varying concentrations of ThDP at pH 8.0 for 8 h was adopted to facilitate the catalysis. The concentration of ThDP affects 6ACA production (Fig. 5B). Without ThDP addition, no 6ACA was produced in this multi-enzyme cascade system. When 0.1 mM ThDP was added, the titer of 6ACA was 20.64 mg/L. Notably, the concentration of 6ACA increased markedly to 46.96 mg/L at 0.5 mM ThDP addition. Consequently, the 0.5 mM of ThDP was set as the optimal dosage.

### 3.4 The confirmation of the rate-limiting enzyme in this artificial iterative cycle

In our initially assembled artificial iterative cycle, 6ACA titer was low (6.68 mg/L), as seen in Fig. 3A. To further improve the titer, a titration experiment was performed with different concentrations of enzymes. We varied the concentration of RaiP, LeuB, LeuC, LeuD, KivD and PadA from 1.0 to 10.0 μM, while the concentration of LeuA* was varied from 1.0 to 20.0 μM. Concentration of 1.0 μM was chosen for the other enzymes. No significant change in the titer of 6ACA was observed with the increased concentration of RaiP. Increasing the concentrations of LeuA^#^to a proper range led to a 3-5 folds increase (6.68 mg/L to 30.18 mg/L) in the titer of 6ACA. When the concentration of LeuA^#^was 20.0 µM, 6ACA reached the highest concentration of 30.18 mg/L shown in Fig. 4. This result suggests that LeuA^#^is a rate-limiting enzyme in the system, consistent with the previous findings from the iterative nonnatural alcohol system (Zhang et al., 2008). Increasing concentrations of LeuB, LeuC, LeuD, KivD and PadA within a specific range resulted in only a modest increase in the titer of 6ACA. No further titer improvement was observed when the enzyme concentrations reached 2.0 μM for LeuC, LeuD and PadA, 4.0 μM for LeuB, whereas 5.0 μM of KivD inhibited 6ACA production. The optimal molar ratio of RaiP: LeuA^#^:LeuB: LeuC: LeuD: KivD: PadA was determined as 1:20:4:2:2:5:2 in this artificial iterative pathway, which was inferred from the titration studies, as seen in Fig. 4.

**Fig. 4.**
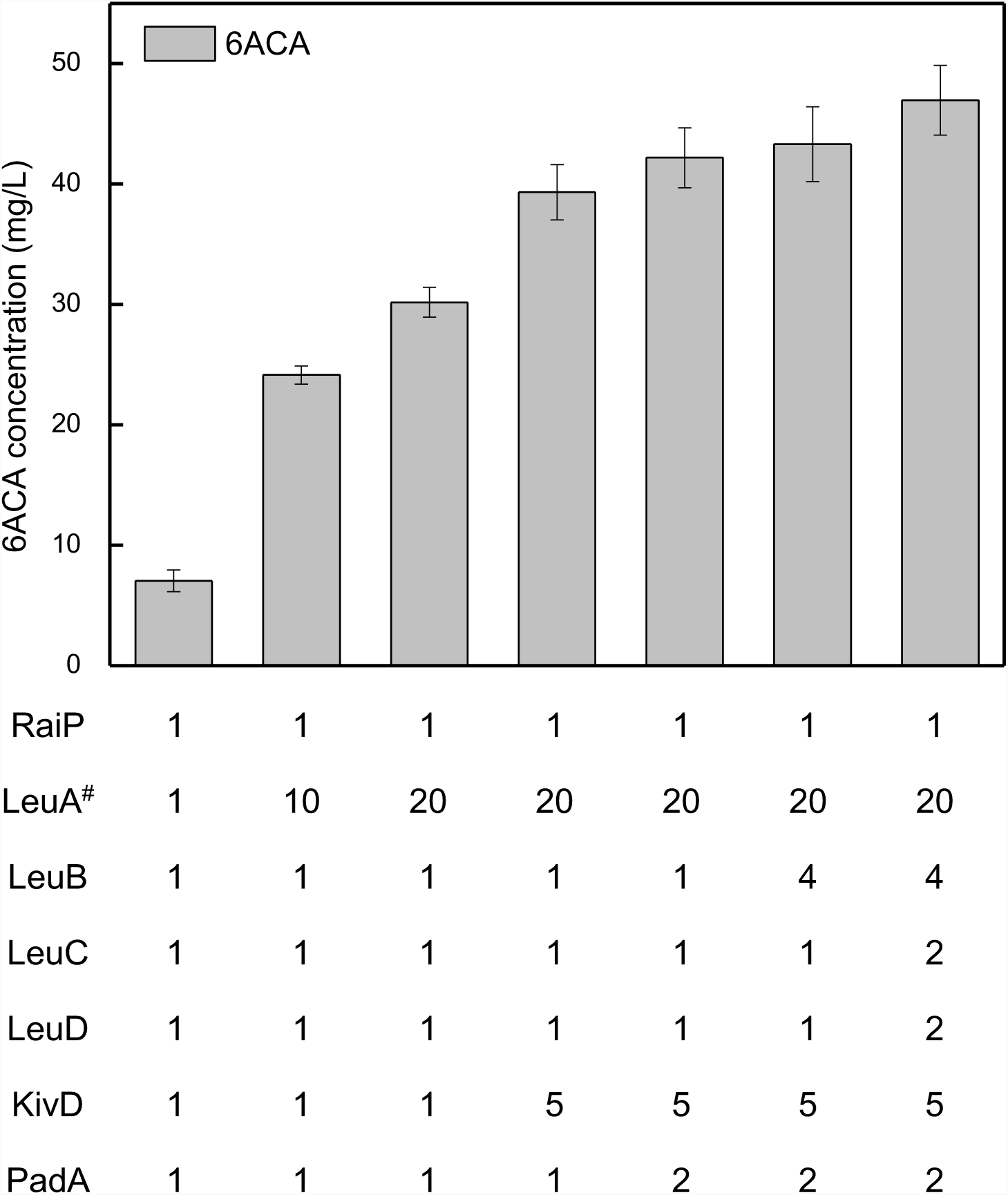
The optimal molar ratio of RaiP:LeuA^#^:LeuB:LeuC:LeuD:KivD:PadA for the production of NNSCAAs. NNSCAAs, Nonnatural straight chain amino acids. 6ACA, 6-Aminocaproate. All experiments were performed a minimum of three independent sets. All error bars represent standard deviations with n ≥ 3 independent reactions.

**Fig. 5.**
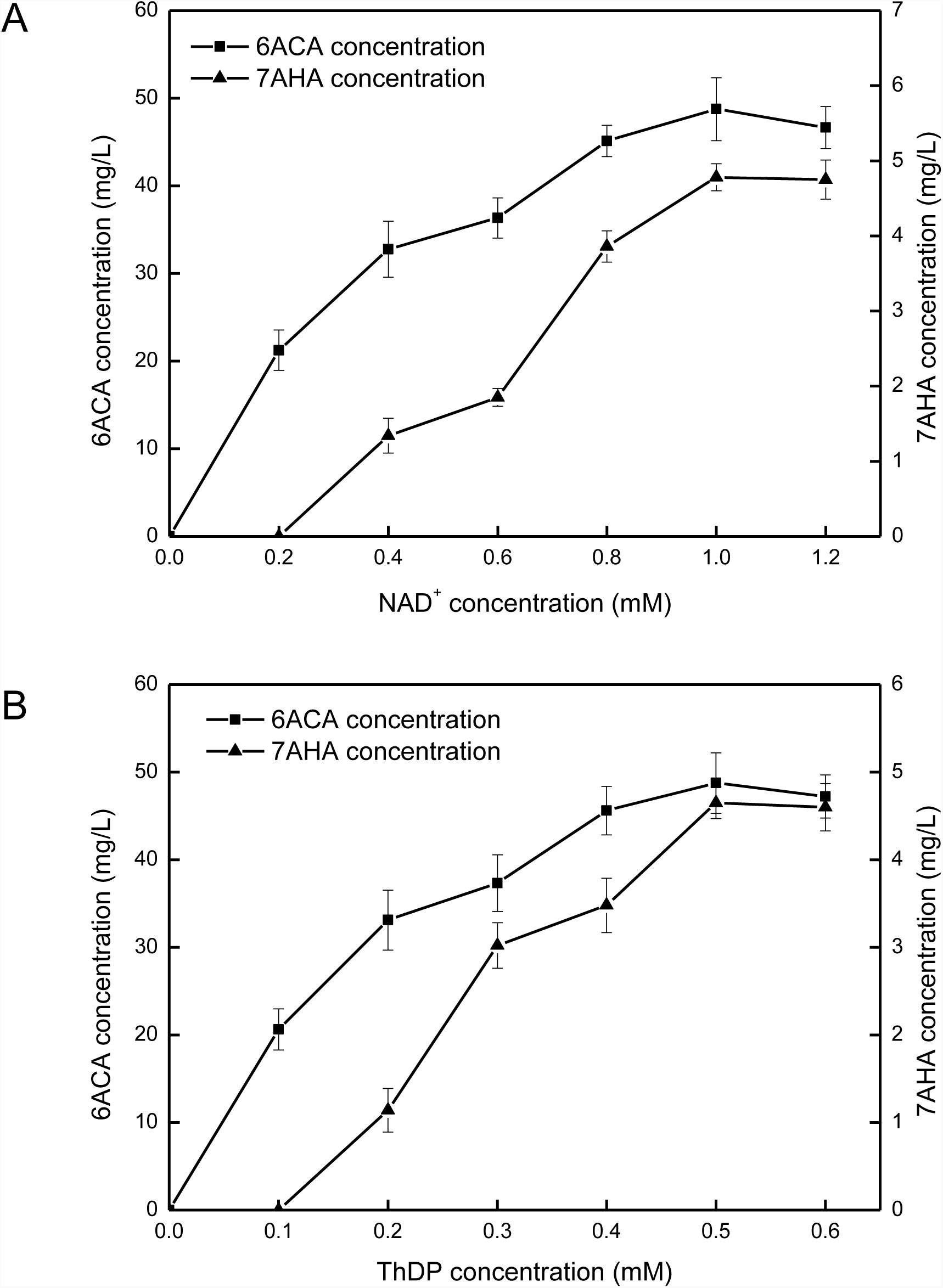
Coenzymes optimizations lead to increased yields of NNSCAAs production. A: Reactions for NNSCAAs production from L-lysine were performed using different sets of NAD^+^. Each assay mixture included 2.0 mM acetyl-CoA, 1.0 μM RaiP, 20.0 μM LeuA^#^, 4.0 μM LeuB, 2.0 μM LeuC, 2.0 μM LeuD, 5.0 μM KivD, 2.0 μM PadA, 2.5 mM L-lysine, 1.0 mM MgCl_2_, 1.0 mM TCEP, 0.5 mM ThDP, and varying concentrations of NAD^+^. Reactions incubated for 8 h at 37 °C. B: Reactions for NNSCAAs production from L-lysine were performed using different sets of ThDP. Each assay mixture included 2.0 mM acetyl-CoA, 1.0 μM RaiP, 20.0 μM LeuA^#^, 4.0 μM LeuB, 2.0 μM LeuC, 2.0 μM LeuD, 5.0 μM KivD, 2.0 μM PadA, 2.5 mM L-lysine, 1.0 mM MgCl_2_, 1.0 mM TCEP, 1.0 mM ThDP, and varying concentrations of ThDP. Reactions incubated for 8 h at 37 °C. 6ACA, 6-Aminocaproate. 7AHA, 7-Aminoheptanoate. All experiments were performed a minimum of three independent sets. All error bars represent standard deviations with n ≥ 3 independent reactions.

**Fig. 6.**
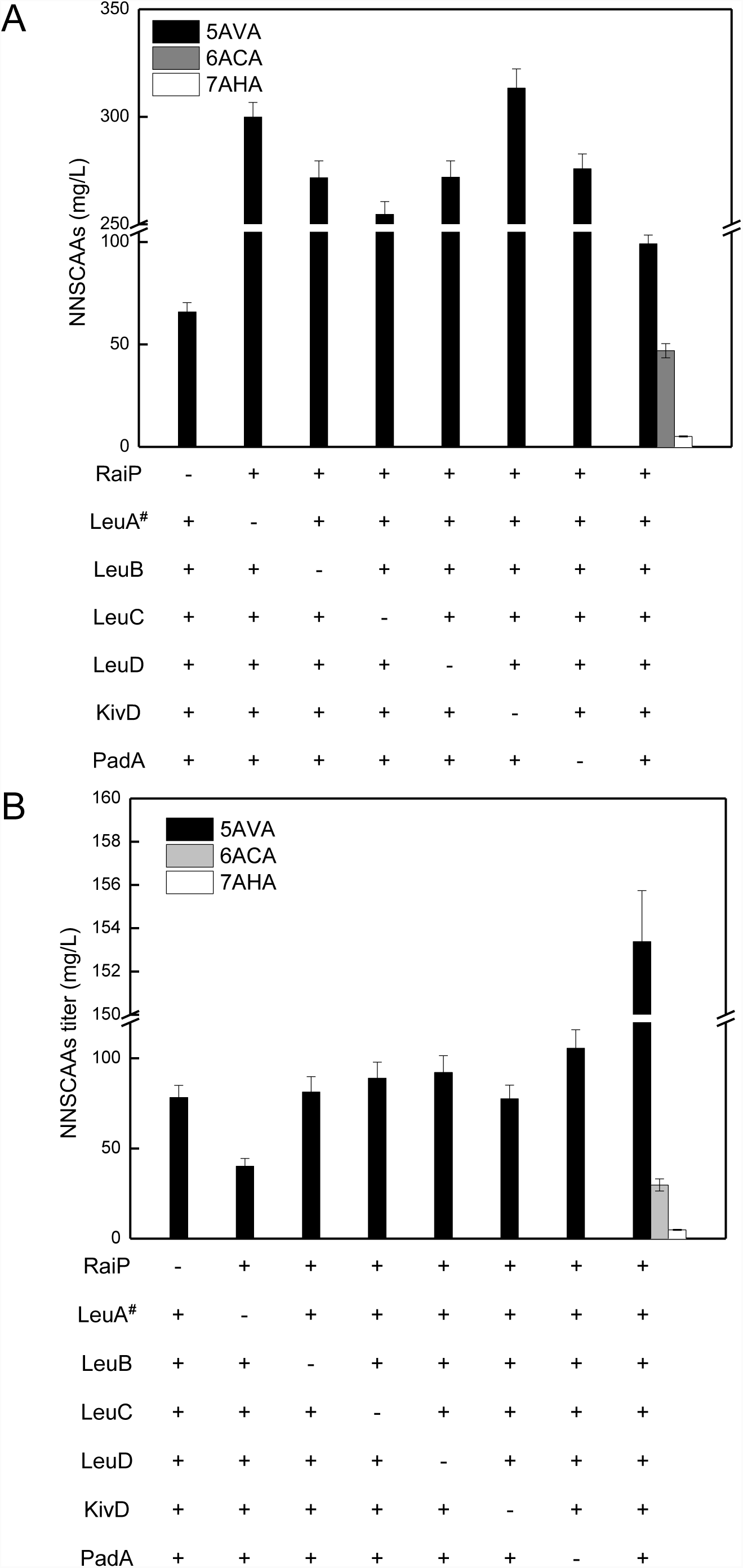
Biosynthesis of NNSCAAs achieved via an artificial iterative carbon-chain-extension cycle. A: Iterative carbon-chain-extension cycle reactions for NNSCAAs production from L-lysine were carried out using seven purified enzymes mixed together (1:20:4:2:2:5:2 based on purified protein quantification) with 1.0 mM MgCl_2_, 1.0 mM TCEP, 50 mM KPB, and coenzymes (ThDP, NAD^+^). These purified enzymes individually contained RaiP, LeuA^#^, LeuB, LeuC, LeuD, KivD and PadA selectively overexpressed at 20 °C. B: RCCEC reactions for NNSCAAs production from L-lysine were carried out using seven crude lysates mixed together (1:20:4:2:2:5:2 based on total protein quantification) with 1.0 mM MgCl_2_, 1.0 mM TCEP, 50 mM KPB, and coenzymes (ThDP, NAD^+^). These lysates individually contained RaiP, LeuA^#^, LeuB, LeuC, LeuD, KivD and PadA selectively overexpressed at 20 °C. NNSCAAs, Nonnatural straight chain amino acids. 5AVA, 5-Aminovalerate. 6ACA, 6-Aminocaproate. 7AHA, 7-Aminoheptanoate. All experiments were performed a minimum of three independent sets. All error bars represent standard deviations with n ≥ 3 independent reactions.

### 3.5 Assembling a nonnatural NNSCAA biosynthetic pathway in *E. coli*

To the best of our knowledge, 6ACA is a nonnatural specialty chemical that could not be directly biosynthesized by any microorganism naturally. Based on the enzyme catalysis result obtained *in vitro*, we sought to build this artificial iterative cycle metabolic pathway in *E. coli* BL21 (DE3) to produce 6ACA by expressing seven enzymes (RaiP, LeuA^#^, LeuB, LeuC, LeuD, KivD and PadA), as seen in Fig. 1. The resulted engineered *E. coli* strain produces total NNSCAAs at a titer of 2.18 g/L from L-lysine. After 36 h of aerobic cultivation, there was no accumulation of 6ACA in the control strain. Whereas the engineered strain produced 6ACA with a peak concentration of 24.12 mg/L at 36 h shown in Fig. 7. The successful assembly of this iterative pathway in *E. coli* validates the *in vitro* pathway design for producing L-lysine-derived NNSCAAs.

**Fig. 7.**
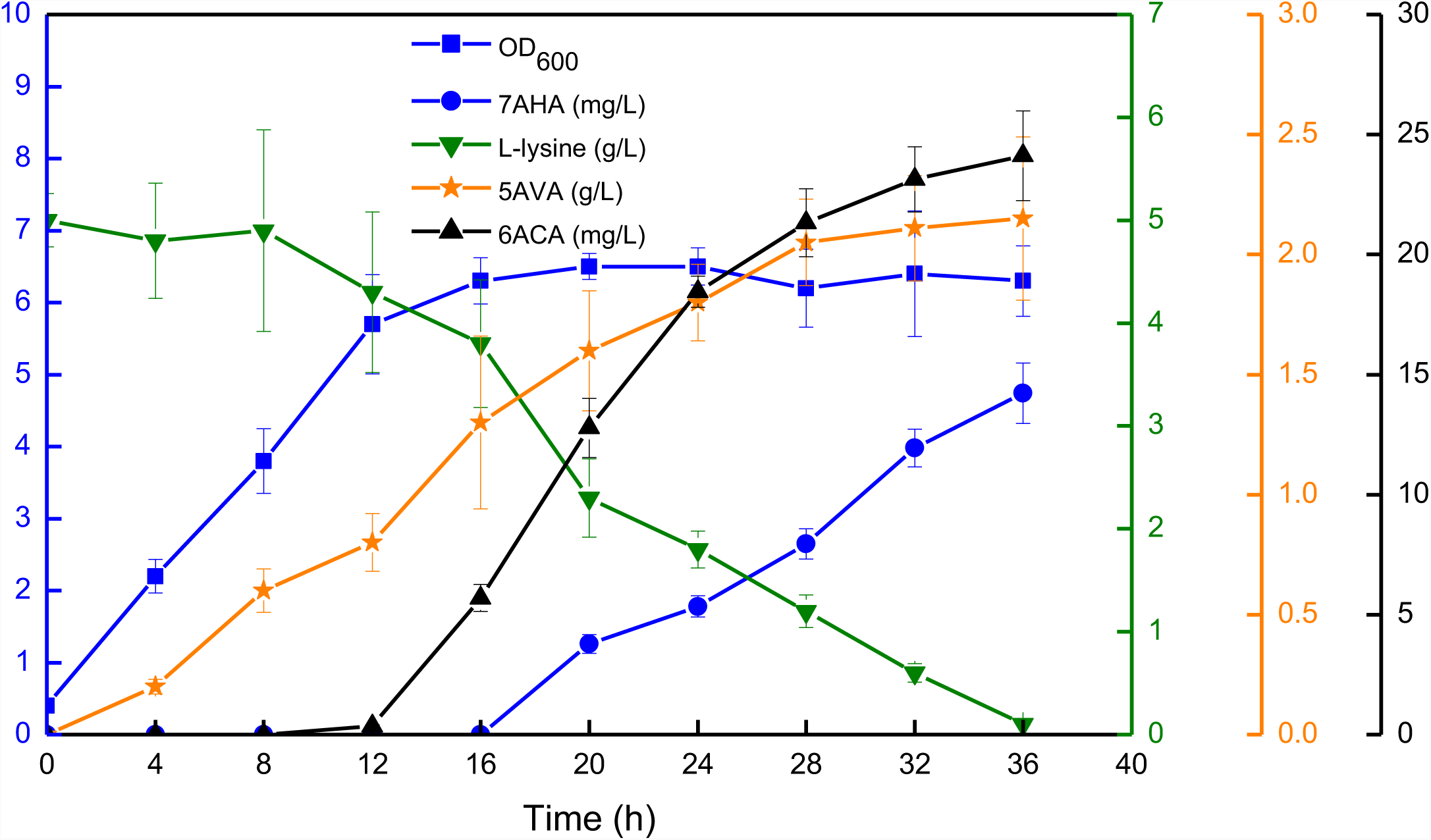
NNSCAAs synthesis by engineered strain CJ04 in 250 mL flask. The cells were grown in 40 mL LB supplemented with 100 μg/mL ampicillin, 50 μg/mL kanamycin, 0.5 mM of IPTG, 5 g/L L-lysine, 1.0 mM MgSO_4_and 0.5 mM ThDP at 37 °C with 250 RPM orbital shaking. Strain CJ04 is strain BL21(DE3) plus plasmids pIVC4 and pET21aRKP. NNSCAAs, Nonnatural straight chain amino acids. 5AVA, 5-Aminovalerate. 6ACA, 6-Aminocaproate. 7AHA, 7-Aminoheptanoate. All experiments were performed a minimum of three independent sets. All error bars represent standard deviations with n ≥ 3 independent reactions.

Metabolic engineering of L-leucine biosynthesis using LeuABCD has been thoroughly explored in *E. coli* (Shen and Liao, 2008; Xiong et al., 2012). Further exploitation of LeuA for the production of longer-chain α-ketoacid and alcohols from branched-chain amino acids was also demonstrated recently (Zhang et al., 2008). The resulting strains produced 6.5 mg/L of 3-methyl-1-pentanol, 17.4 mg/L of 1-hexanol and 22.0 mg/L of 5-methyl-1-heptanol, respectively (Zhang et al., 2008). Connor *et al.* showed that 800 mg/L of 3-methyl-1-butanol could be produced via carbon-chain-extension pathway (Connor and Liao, 2008). Atsumi *et al.* reported that 44.4 mg/L of 1-butanol could be achieved via carbon-chain-extension pathway (Atsumi et al., 2008). In this work, we utilize the L-lysine-derivative 2-keto-6-aminocaproate as substrate for LeuA^#^BCD, therefore broadening the known substrate profile to include terminal-amino-group-containing α-ketoacids. Metabolic engineering of 6ACA production was previously attempted in the strain eAKP-744 (Turk et al., 2016). However, the reported strategy was bottlenecked at the transamination step (Turk et al., 2016; Zhang et al., 2010). The production of 6ACA in the artificial iterative cycle developed in our study circumvents the transamination step, therefore greatly improves the efficiency of the whole process. We anticipate that the efficiency of this NNSCAA biosynthetic pathway can be further improved by optimizing various components of the system.

The synthetic “+1” carbon-chain-extension pathway with α-ketoacids as substrates has been widely exploited to produce chain-elongated alcohols and acids (Atsumi et al., 2008; Marcheschi et al., 2012; Zhang et al., 2008). The carbon-carbon condensation reaction catalyzed by LeuA follows the Felkin-Anh model for nucleophilic attack on a carbonyl (Benjamin and Collins, 1973; Cherest et al., 1968; Marcheschi et al., 2012). However, previous studies were based on α-ketoacids substrates without R groups and the reactions were not iterative. Quantum mechanical calculation predicts that different R groups containing -NH_2_, -SCH_3_, -SOCH_3_, -SH, -COOH, -COH or –OH would not significantly impact the barrier for carbon-chain-extension reaction (Felnagle et al., 2012; Marcheschi et al., 2012). Howell *et al*. have reported that *aksA* in *Methanococcus janaschii* is analogous to *leuA* in *E. coli* (Howell et al., 2000). It is reported that the R group of AksA includes -COOH. α-Ketoglutarate could be converted into α-ketoglutarate, α-ketoglutarate and α-ketoglutarate one after another in *methanoarchaea* by overexpressing AksADEF (Howell et al., 2000; Howell et al., 1998). While R group of LeuA expected here is -NH_2_, the sequence identity between LeuA and AksA is 42%, as seen in Fig. S4. Several key residues at the LeuA active site have been verified to play essential role in controlling the pocket size and hence substrate specifictity, in particular, H97, S139 and G462 shown in Fig. S2 (Chen et al., 2017; Xiong et al., 2012). The natural substrates of LeuA are 2-ketoisovalerate and 2-ketobutyrate (Shen and Liao, 2008). In this study, our engineered *E. coli* strain could use 2-keto-6-aminocaproate as the alternative substrate to simultaneously produce 5AVA, 6ACA and 7AHA from L-lysine with a titer of total at 2.18 g/L, as seen in Fig. 7.

## 4. Conclusion

In summary, we devised a novel strategy to biosynthesize NNSCAAs with different carbon lengths simultaneously in engineered *E. coli*. This work provides a sustainable route for industrial NNSCAA production from renewable feedstocks using metabolic engineering. By coupling new metabolic pathway design and condition optimization, 99.15 mg/L and 46.96 mg/L of 5AVA and 6ACA could be produced *in vitro*, as seen in Fig. 3B and Fig. 6, respectively. Furthermore, production of 4.78 g/L 7AHA could be demonstrated for the first time, which could be used to synthesize new polyurethane nylon 7. LeuA^#^is the rate-limiting enzyme in this artificial iterative reaction, which may be further improved by directed evolution approach in the future. Our success in metabolic engineering the production of 6-aminocaproate and other NNSCAAs via the artificial iterative carbon-chain-extension cycle suggests that similar strategies could be developed to produce other medium-chain-length acids with -NH_2_, -SCH_3_, -SOCH_3_, -SH, -COOH, -COH or -OH functional groups.

### Notes

The authors declare no competing financial interest.

## Acknowledgements

This research was supported by the Fundamental Research Funds for the Central Universities (Project No. 106112017CDJXFLX0014, 2018CDQYHG0010), the Science and Technology Support Program of Tianjin, China (15PTCYSY00020), the Research Equipment Program of Chinese Academy of Sciences (YJKYYQ20170023), the Key Laboratory of Systems Microbial Biotechnology, Tianjin Institute of Industrial Biotechnology, Chinese Academy of Sciences, and the Henan Provincial Science and technology Open cooperation projects (162106000014). This work is also partially supported by Open Funding Project of the State Key Laboratory of Bioreactor Engineering, Shanghai, China.

## Abbreviations

NNSCAA: Nonnatural straight chain amino acid
5AVA: 5-Aminovalerate
6ACA: 6-Aminocaproate
7AHA: 7-Aminoheptanoate
RaiP: L-lysine α oxidase
LeuA: α-Isopropylmalate synthase
LeuA*: LeuA mutants
**LeuA**^**#**^: LeuA with H97L/S139G/G462D mutations
LeuB: 3-Isopropylmalate dehydrogenase
LeuC: 3-Isopropylmalate dehydratase
LeuD: 3-Isopropylmalate dehydratase
KivD: α-Ketoacid decarboxylase
PadA: Aldehyde dehydrogenase
ThDP: Thiamine diphosphate
TCEP: Tris (2-carboxyethyl) phosphine
KPB: Potassium phosphate buffer
LC-MS: Liquid chromatography-mass spectrometry
SDS-PAGE: Sodium dodecyl sulfate polyacrylamide gel electrophoresis
4AAP: 4-Aminoantipyrine

