## Supplemental materials for "Production of nonnatural straight-chain amino acid 6-aminocaproate via an artificial iterative carbon-chain-extension cycle"

**Supplementary Table S1. *Escherichia coli* strains and plasmids used in this study.**

|  | Relevant genotype or description | Sources |
| --- | --- | --- |
| *Escherichia coli* strains |  |  |
| DH5α | Wild type | Novagen |
| BL21(DE3) | Wild type | (Cheng et al., 2018a) |
| CJ03 | BL21(DE3) harboring pIVC3 and pETaRKP | This study |
| CJ04 | BL21(DE3) harboring pIVC4 and pETaRKP | This study |
| Plasmids |  |  |
| pET21a | Empty plasmid used as control, Amp^R^ | (Cheng et al., 2018b) |
| pZA22 | Empty plasmid used as control, Kan^R^ | (Xiong et al., 2012) |
| pET22b | Empty plasmid used as control, Amp^R^ | This study |
| pCJ01 | pET21a-*raip*, pET21a carries a L-lysine α-oxidase (*raiP*) gene from *Scomber japonicas* with *Nde*I and *BamH*I restrictions, Amp^R^ | (Cheng et al., 2018b) |
| pETaraiP | pET21a-*raip*, pET21a carries a L-lysine α-oxidase (*raiP*) gene from *Scomber japonicas* with *Nde*I and *Xho*I restrictions, Amp^R^ | This study |
| pETaleuA | pET21a-*leuA*, pET21a carries a α-isopropylmalate synthase (*leuA*) gene from *Escherichia coli*, Amp^R^ | This study |
| pETaleuB | pET21a-*leuB*, pET21a carries a 3-isopropylmalate dehydrogenase (*leuB*) gene from *Escherichia coli*, Amp^R^ | This study |
| pETaleuC | pET21a-*leuC*, pET21a carries a 3-isopropylmalate dehydratase large subunit (*leuC*) gene from *Escherichia coli*, Amp^R^ | This study |
| pETaleuD | pET21a-*leuD*, pET21a carries a 3-isopropylmalate dehydratase small subunit (*leuD*) gene from *Escherichia coli*, Amp^R^ | This study |
| pETbkivD | pET22b-*kivD*, pET22b carries a α-ketoacid decarboxylase (*kivD*) gene from *Lactococcus lactis*, Amp^R^ | This study |
| pETapadA | pET21a-*padA*, pET21a carries a aldehyde dehydrogenase (*padA*) gene from *Escherichia coli*, Amp^R^ | This study |
| pIVC3 | pZA22-*leuA*-*leuB*-*leuC*-*leuD*, pZA22 carries a α-isopropylmalate synthase (*leuA*) gene, a 3-isopropylmalate dehydrogenase (*leuB*) gene, a 3-isopropylmalate dehydratase large subunit (*leuC*) gene and a 3-isopropylmalate dehydratase small subunit (*leuD*) gene from *Escherichia coli*, Kan^R^ | This study |
| pIVC4 | pZA22-*leuA**-*leuB*-*leuC*-*leuD*, pZA22 carries a α-isopropylmalate synthase mutant (H97L/S139G/G462D) gene, a 3-isopropylmalate dehydrogenase (*leuB*) gene, a 3-isopropylmalate dehydratase large subunit (*leuC*) gene and a 3-isopropylmalate dehydratase small subunit (*leuD*) gene from *Escherichia coli*, Kan^R^ | This study |
| pETaRKP | pET21a-*raiP*-*kivD*-*padA*, pET21a carries a L-lysine α-oxidase (*raiP*) gene from *Scomber japonicas*, a α-ketoacid decarboxylase (*kivD*) gene from *Lactococcus lactis* and a aldehyde dehydrogenase (*padA*) gene from *Escherichia coli*, Amp^R^ | This study |
| pETaleuA*(G462D) | pET21a carries a α-isopropylmalate synthase mutant (G462D) gene from *Escherichia coli*, Amp^R^ | This study |
| pETaleuA*( H97A/G462D) | pET21a carries a α-isopropylmalate synthase mutant (H97A/G462D) gene from *Escherichia coli*, Amp^R^ | This study |
| pETaleuA*( H97C/G462D) | pET21a carries a 2-isopropylmalate synthase mutant (H97C/G462D) gene from *Escherichia coli*, Amp^R^ | This study |
| pETaleuA*( H97D/G462D) | pET21a carries a 2-isopropylmalate synthase mutant (H97D/G462D) gene from *Escherichia coli*, Amp^R^ | This study |
| pETaleuA*( H97E/G462D) | pET21a carries a 2-isopropylmalate synthase mutant (H97E/G462D) gene from *Escherichia coli*, Amp^R^ | This study |
| pETaleuA*( H97F/G462D) | pET21a carries a 2-isopropylmalate synthase mutant (H97F/G462D) gene from *Escherichia coli*, Amp^R^ | This study |
| pETaleuA*( H97G/G462D) | pET21a carries a 2-isopropylmalate synthase mutant (H97G/G462D) gene from *Escherichia coli*, Amp^R^ | This study |
| pETaleuA*( H97I/G462D) | pET21a carries a 2-isopropylmalate synthase mutant (H97I/G462D) gene from *Escherichia coli*, Amp^R^ | This study |
| pETaleuA*( H97K/G462D) | pET21a carries a 2-isopropylmalate synthase mutant (H97K/G462D) gene from *Escherichia coli*, Amp^R^ | This study |
| pETaleuA*( H97L/G462D) | pET21a carries a 2-isopropylmalate synthase mutant (H97L/G462D) gene from *Escherichia coli*, Amp^R^ | This study |
| pETaleuA*( H97M/G462D) | pET21a carries a 2-isopropylmalate synthase mutant (H97M/G462D) gene from *Escherichia coli*, Amp^R^ | This study |
| pETaleuA*( H97N/G462D) | pET21a carries a 2-isopropylmalate synthase mutant (H97N/G462D) gene from *Escherichia coli*, Amp^R^ | This study |
| pETaleuA*( H97P/G462D) | pET21a carries a 2-isopropylmalate synthase mutant (H97P/G462D) gene from *Escherichia coli*, Amp^R^ | This study |
| pETaleuA*( H97Q/G462D) | pET21a carries a 2-isopropylmalate synthase mutant (H97Q/G462D) gene from *Escherichia coli*, Amp^R^ | This study |
| pETaleuA*( H97R/G462D) | pET21a carries a 2-isopropylmalate synthase mutant (H97R/G462D) gene from *Escherichia coli*, Amp^R^ | This study |
| pETaleuA*( H97S/G462D) | pET21a carries a 2-isopropylmalate synthase mutant (H97S/G462D) gene from *Escherichia coli*, Amp^R^ | This study |
| pETaleuA*( H97T/G462D) | pET21a carries a 2-isopropylmalate synthase mutant (H97T/G462D) gene from *Escherichia coli*, Amp^R^ | This study |
| pETaleuA*( H97V/G462D) | pET21a carries a 2-isopropylmalate synthase mutant (H97V/G462D) gene from *Escherichia coli*, Amp^R^ | This study |
| pETaleuA*( H97W/G462D) | pET21a carries a 2-isopropylmalate synthase mutant (H97W/G462D) gene from *Escherichia coli*, Amp^R^ | This study |
| pETaleuA*( H97Y/G462D) | pET21a carries a 2-isopropylmalate synthase mutant (H97Y/G462D) gene from *Escherichia coli*, Amp^R^ | This study |
| pETaleuA*( S139A/G462D) | pET21a carries a 2-isopropylmalate synthase mutant (S139A/G462D) gene from *Escherichia coli*, Amp^R^ | This study |
| pETaleuA*( S139C/G462D) | pET21a carries a 2-isopropylmalate synthase mutant (S139C/G462D) gene from *Escherichia coli*, Amp^R^ | This study |
| pETaleuA*( S139D/G462D) | pET21a carries a 2-isopropylmalate synthase mutant (S139D/G462D) gene from *Escherichia coli*, Amp^R^ | This study |
| pETaleuA*( S139E/G462D) | pET21a carries a 2-isopropylmalate synthase mutant (S139E/G462D) gene from *Escherichia coli*, Amp^R^ | This study |
| pETaleuA*( S139F/G462D) | pET21a carries a 2-isopropylmalate synthase mutant (S139F/G462D) gene from *Escherichia coli*, Amp^R^ | This study |
| pETaleuA*( S139G/G462D) | pET21a carries a 2-isopropylmalate synthase mutant (S139G/G462D) gene from *Escherichia coli*, Amp^R^ | This study |
| pETaleuA*( S139H/G462D) | pET21a carries a 2-isopropylmalate synthase mutant (S139H/G462D) gene from *Escherichia coli*, Amp^R^ | This study |
| pETaleuA*( S139I/G462D) | pET21a carries a 2-isopropylmalate synthase mutant (S139I/G462D) gene from *Escherichia coli*, Amp^R^ | This study |
| pETaleuA*( S139K/G462D) | pET21a carries a 2-isopropylmalate synthase mutant (S139K/G462D) gene from *Escherichia coli*, Amp^R^ | This study |
| pETaleuA*( S139L/G462D) | pET21a carries a 2-isopropylmalate synthase mutant (S139L/G462D) gene from *Escherichia coli*, Amp^R^ | This study |
| pETaleuA*( S139M/G462D) | pET21a carries a 2-isopropylmalate synthase mutant (S139M/G462D) gene from *Escherichia coli*, Amp^R^ | This study |
| pETaleuA*( S139N/G462D) | pET21a carries a 2-isopropylmalate synthase mutant (S139N/G462D) gene from *Escherichia coli*, Amp^R^ | This study |
| pETaleuA*( S139P/G462D) | pET21a carries a 2-isopropylmalate synthase mutant (S139P/G462D) gene from *Escherichia coli*, Amp^R^ | This study |
| pETaleuA*( S139Q/G462D) | pET21a carries a 2-isopropylmalate synthase mutant (S139Q/G462D) gene from *Escherichia coli*, Amp^R^ | This study |
| pETaleuA*( S139R/G462D) | pET21a carries a 2-isopropylmalate synthase mutant (S139R/G462D) gene from *Escherichia coli*, Amp^R^ | This study |
| pETaleuA*( S139T/G462D) | pET21a carries a 2-isopropylmalate synthase mutant (S139T/G462D) gene from *Escherichia coli*, Amp^R^ | This study |
| pETaleuA*( S139V/G462D) | pET21a carries a 2-isopropylmalate synthase mutant (S139V/G462D) gene from *Escherichia coli*, Amp^R^ | This study |
| pETaleuA*( S139W/G462D) | pET21a carries a 2-isopropylmalate synthase mutant (S139W/G462D) gene from *Escherichia coli*, Amp^R^ | This study |
| pETaleuA*( S139Y/G462D) | pET21a carries a 2-isopropylmalate synthase mutant (S139Y/G462D) gene from *Escherichia coli*, Amp^R^ | This study |
| pETaleuA*( H97A/S139G/G462D) | pET21a carries a 2-isopropylmalate synthase mutant (H97A/S139Y/G462D) gene from *Escherichia coli*, Amp^R^ | This study |
| pETaleuA*( H97A/S139I/G462D) | pET21a carries a 2-isopropylmalate synthase mutant (H97A/S139I/G462D) gene from *Escherichia coli*, Amp^R^ | This study |
| pETaleuA*( H97G/S139G/G462D) | pET21a carries a 2-isopropylmalate synthase mutant (H97G/S139G/G462D) gene from *Escherichia coli*, Amp^R^ | This study |
| pETaleuA*( H97G/S139I/G462D) | pET21a carries a 2-isopropylmalate synthase mutant (H97G/S139I/G462D) gene from *Escherichia coli*, Amp^R^ | This study |
| pETaleuA*( H97L/S139I/G462D) | pET21a carries a 2-isopropylmalate synthase mutant (H97L/S139I/G462D) gene from *Escherichia coli*, Amp^R^ | This study |
| pETaleuA*( H97L/S139G/G462D) | pET21a-*leuA**, pET21a carries a 2-isopropylmalate synthase mutant (H97L/S139G/G462D) gene from *Escherichia coli*, Amp^R^ | This study |

**Supplementary Table S2. List of primers used in this study.**

| Primers names | Nucleotide sequence (5’-3’) |
| --- | --- |
| pETaraiP-F | 5′-GGAATTCCATATGGAGCACCTGGCAGACTGTCTGG-3′ |
| pETaraiP-R | 5′-CCGCTCGAGCAGTTCGTCCTTGGTATG-3′ |
| pETaleuA-F | 5′-GGAATTCCATATGAGCCAGCAAGTG-3′ |
| pETaleuA-R | 5′-CCGCTCGAGAACGGTTTCTTTGTTA-3′ |
| pETaleuB-F | 5′-CGGGATCCATGTCGAAGAATTACC-3′ |
| pETaleuB-R | 5′-CCGCTCGAGCACCCCTTCTGCTACATAG-3′ |
| pETaleuC-F | 5′-GGAATTCCATATGGCTAAGACGTTATACG-3′ |
| pETaleuC-R | 5′-CCGCTCGAGTTTAATGTTGCGAATGTCGGC-3′ |
| pETaleuD-F | 5′-GGAATTCCATATGGCAGAGAAATTTATC-3′ |
| pETaleuD-R | 5′-CCGCTCGAGATTCATAAACGCAGGTTG-3′ |
| pETapadA-F | 5′-GGAATTCCATATGACAGAGCCGCATGTAGC-3′ |
| pETapadA-R | 5′-CCGCTCGAGATACCGTACACACACC-3′ |
| T7 promoter | 5′-TAATACGACTCACTATAGG-3′ |
| T7 terminator | 5′-GCTAGTTATTGCTCAGCGG-3′ |
| G462D-F | GACCAGGTGGACATCGTTGCC |
| G462D-R | CAGGGCATCTTTGCCATGGC |
| H97A-F | GCTACCTTCATCGCCACCAGTC |
| H97C-F | TGCACCTTCATCGCCACCAGTC |
| H97D-F | GACACCTTCATCGCCACCAGTC |
| H97E-F | GAGACCTTCATCGCCACCAGTC |
| H97F-F | TTCACCTTCATCGCCACCAGTC |
| H97G-F | GGAACCTTCATCGCCACCAGTC |
| H97I-F | ATCACCTTCATCGCCACCAGTC |
| H97K-F | AAGACCTTCATCGCCACCAGTC |
| H97L-F | CTGACCTTCATCGCCACCAGTC |
| H97M-F | ATGACCTTCATCGCCACCAGTC |
| H97N-F | AACACCTTCATCGCCACCAGTC |
| H97P-F | CCTACCTTCATCGCCACCAGTC |
| H97Q-F | ACGACCTTCATCGCCACCAGTC |
| H97R-F | CGTACCTTCATCGCCACCAGTC |
| H97S-F | AGTACCTTCATCGCCACCAGTC |
| H97T-F | ACGACCTTCATCGCCACCAGTC |
| H97V-F | GTCACCTTCATCGCCACCAGTC |
| H97W-F | TGGACCTTCATCGCCACCAGTC |
| H97Y-F | TACACCTTCATCGCCACCAGTC |
| H97-R | GATGCGAAATGCCTCTGCAAC |
| S139A-F | GCTTGCGAAGATGCCGGCCGC |
| S139C-F | TGCTGCGAAGATGCCGGCCGC |
| S139D-F | GACTGCGAAGATGCCGGCCGC |
| S139E-F | GAGTGCGAAGATGCCGGCCGC |
| S139F-F | TTCTGCGAAGATGCCGGCCGC |
| S139G-F | GGCTGCGAAGATGCCGGCCGC |
| S139H-F | CATTGCGAAGATGCCGGCCGC |
| S139I-F | ATCTGCGAAGATGCCGGCCGC |
| S139K-F | AAGTGCGAAGATGCCGGCCGC |
| S139L-F | CTGTGCGAAGATGCCGGCCGC |
| S139M-F | ATGTGCGAAGATGCCGGCCGC |
| S139N-F | AACTGCGAAGATGCCGGCCGC |
| S139P-F | CCTTGCGAAGATGCCGGCCGC |
| S139Q-F | CAGTGCGAAGATGCCGGCCGC |
| S139R-F | CGTTGCGAAGATGCCGGCCGC |
| S139T-F | ACGTGCGAAGATGCCGGCCGC |
| S139V-F | TGCTGCGAAGATGCCGGCCGC |
| S139W-F | TGGTGCGAAGATGCCGGCCGC |
| S139Y-F | TACTGCGAAGATGCCGGCCGC |
| S139-R | GAACTCAACGTCATCGGTATAG |

Restriction sites are underlined.

**Supplementary Table S3. Gradient elution program for LC and LC-MS.**

| **Time (min)** | **Buffer A (%)** | **Buffer B (%)** |
| --- | --- | --- |
| 0.0 | 3 | 97 |
| 1.0 | 3 | 97 |
| 20.0 | 30 | 70 |
| 30.0 | 35 | 65 |
| 40.0 | 90 | 10 |
| 41.0 | 100 | 0 |
| 45.0 | 100 | 0 |
| 46.0 | 3 | 97 |
| 55.0 | 3 | 97 |

Captions to Supplementary Figures

### Fig. S1. A: Via SDS-PAGE, the gel verifies the selective overexpression of pathway enzymes in *E. coli* BL21(DE3) curde cell lysates. RaiP, L-lysine α-oxidase; LeuA, α-Isopropylmalate synthase; LeuB, 3-isopropylmalate dehydrogenase; LeuC, 3-isopropylmalate dehydratase large subunit; LeuD, 3-isopropylmalate dehydratase small subunit; KivD, α-ketoacid decarboxylase; PadA, aldehyde dehydrogenase. Protein samples were separated by 12% SDS-PAGE and stained with coomassie brilliant blue. Lane 1, Molecular weight markers (kDa). Lane 2, pET21a induced control. Lane 3, RaiP induced. Lane 4, LeuA induced. Lane 5, LeuB induced. Lane 6, LeuC induced. Lane 7, LeuD induced in. Lane 8, KivD induced. Lane 9, PadA induced. The expression was induced with 0.5 mM IPTG. B: Via SDS-PAGE, the gel verifies the selective overexpression of pathway enzymes in *E. coli* BL21(DE3) purified enzymes. Lane 1, Molecular weight marker (kDa). Lane 2, Purification of RaiP. Lane 3, Purification of LeuA. Lane 4, Purification of LeuB. Lane 5, Purification of LeuC. Lane 6, Purification of LeuD. Lane 7, Purification of KivD. Lane 8, Purification of PadA. Proteins were induced with 0.5 mM IPTG and incubated in 20 °C for 16 h.

### Fig. S2. Residues in the active site of LeuA. Binding pocket of *E. coli* LeuA (PDB: 1SR9) complexed with its substrate 2-keto-6-aminocaproate. The active pocket of LeuA which is constituted by a number of hydrophobic residues, including H97, S139, N167 and T171. LeuA, α-isopropylmalate synthase.

### Fig. S3. Via SDS-PAGE, the gel verifies the selective overexpression of various LeuA mutants. Various saturation mutants of LeuA were conducted on His97 and Ser139. Strains were induced with 0.5 mM IPTG and incubated in 20 °C for 16 h.

### Fig. S4. The BLAST result of LeuA (GenBank Accession No. NC_000913) against AksA (GenBank Accession No. NC_021355). The blast result of protein LeuA and AksA was performed by NCBI web server (https://blast.ncbi.nlm.nih.gov/Blast.cgi).

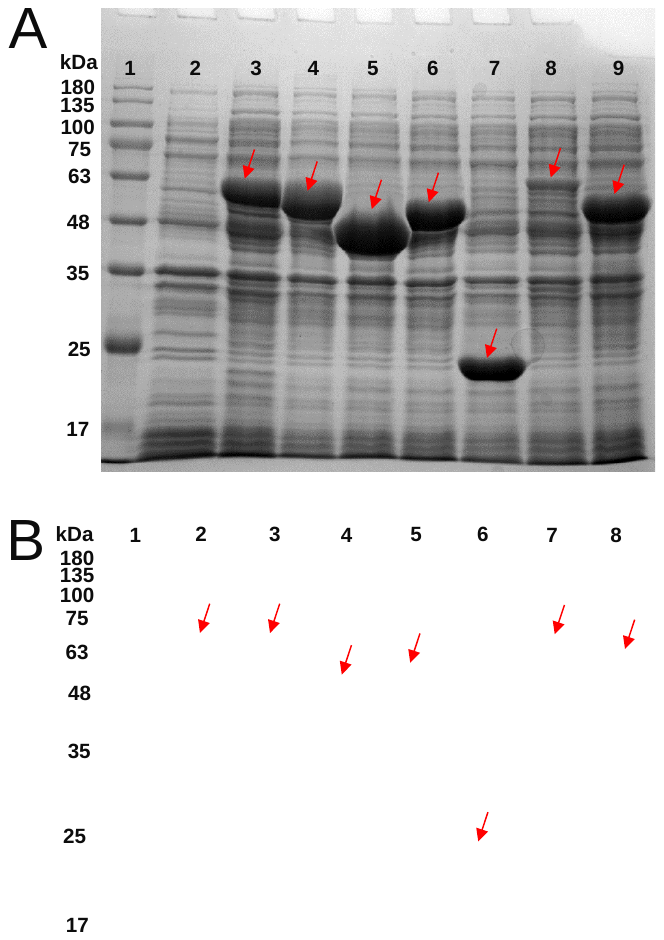

**Fig. S1**

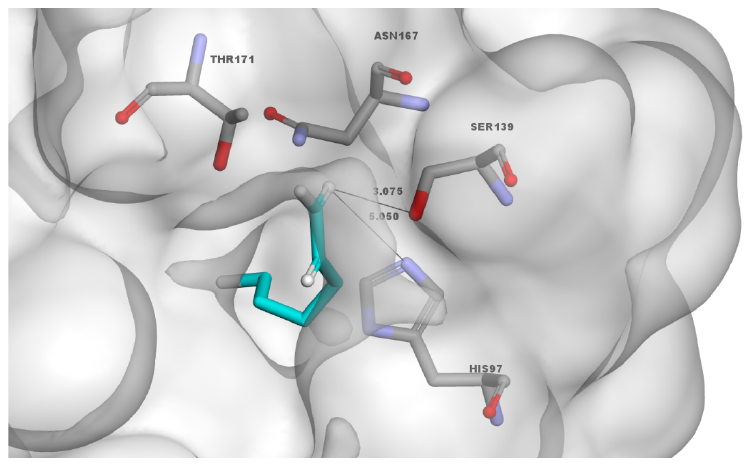

**Fig. S2**

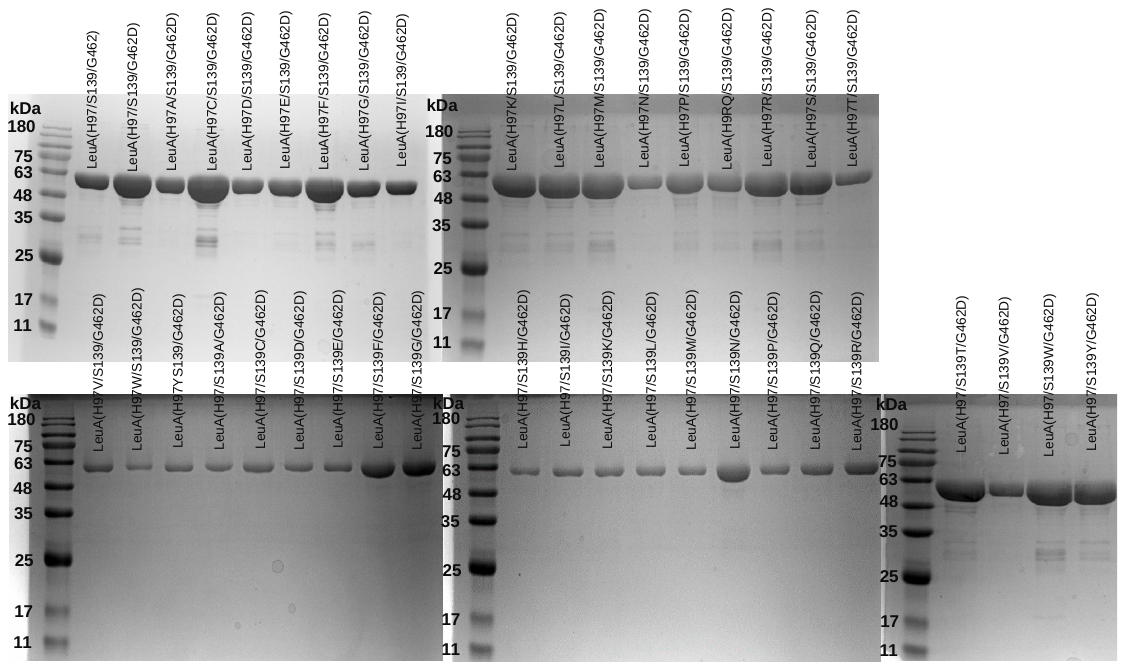

**Fig. S3**

| Score | Expect | Method | Identities | Positives | Gaps |
| --- | --- | --- | --- | --- | --- |
| 234 bits(598) | 1e-76 | Compositional matrix adjust. | 143/344(42%) | 202/344(58%) | 13/344(3%) |

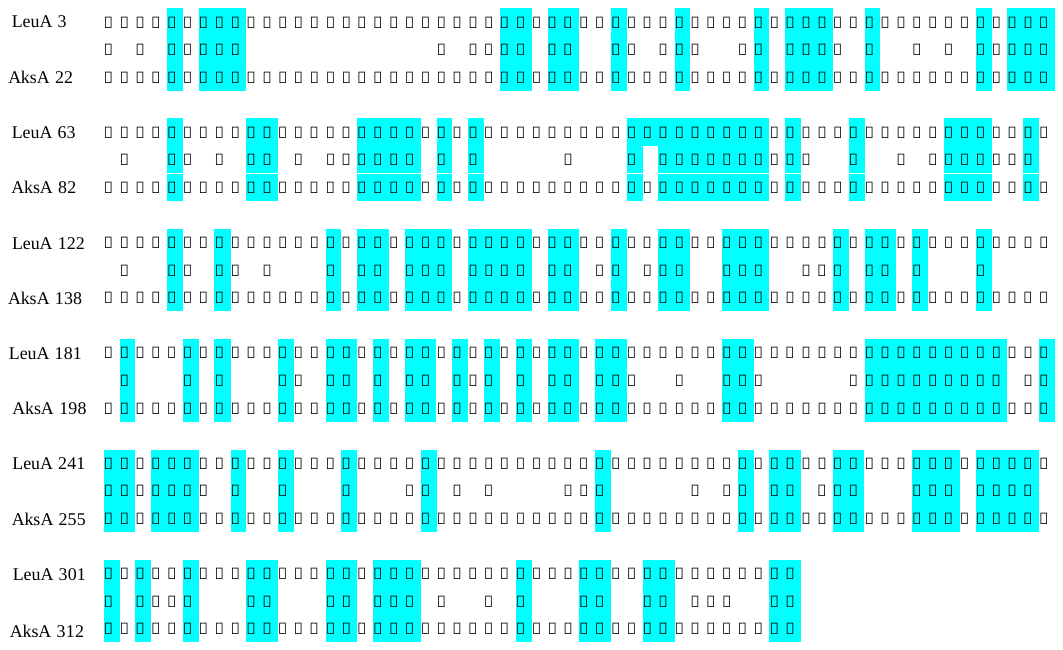

**Fig. S4**
